# Herbivory-induced alterations in cytosolic proteins of pigeon pea (*Cajanus cajan*) leaves

**DOI:** 10.64898/2026.05.07.723431

**Authors:** S Arunima, Prakash Jyoti Kalita, Swapnil Kumar Meshram, Alakesh Das, Rahul Ishwar Patil, Supriya Das, Jagdish Jaba, Debajit Das, Sumita Acharjee

## Abstract

Insect herbivory triggers cytosolic proteome reprogramming by activating defense pathways and modulating key metabolic processes. We found that simulated herbivory in pigeon pea (*Cajanus cajan*) induced reactive oxygen species (ROS) production and molecular alterations within 12 hours (h) of post treatment. We compared the leaf proteome profiles of two cultivated genotypes, ICPL 332 (moderately resistant) and ICPL 87 (susceptible), using two-dimensional polyacrylamide gel electrophoresis (2D-PAGE) coupled with mass spectrometry (MS). More than 220 protein spots were detected in ICPL 332 and over 200 in ICPL 87. Comparative analysis revealed 75 differentially accumulated proteins (DAPs), of which 40 were consistently reproducible across biological replicates. These included 11 unique to ICPL 87, 9 unique to ICPL 332, and 10 common to both genotypes. Among the shared DAPs, ICPL 332 showed five upregulated and five downregulated, whereas ICPL 87 exhibited only two upregulated and eight downregulated. Functional categorization grouped DAPs into primary metabolism, stress response, and growth and development. Proteins related to primary metabolism were largely downregulated in both genotypes, while stress-associated proteins exhibited substantial downregulation in ICPL 87 compared to ICPL 332. Overall, the results demonstrate proteomic adjustments underlying defense responses in pigeon pea genotypes.

## Introduction

Insect pests represent a major biotic constraint to global agriculture, causing yield losses estimated to exceed 18% worldwide [1]. To counteract these threats, plants have evolved sophisticated defense mechanisms, comprising both constitutive adaptations and inducible responses that function across developmental stages. These defense responses are initiated by the recognition of herbivore-associated molecular patterns (HAMPs) present in oral secretions (OS), which contain key molecules such as glucose oxidase, apolipophorin, caspase, scolexin, lectins, phenol oxidase, hemolysin, and damage-associated molecular patterns (DAMPs) released from wounded plant cells [2]. Plant respond to biotic and abiotic stressors by activating complex signaling networks, characterized by calcium fluxes and phosphorylation cascades, which are central to the regulation and coordination defense responses [3].

Herbivore attack also induces extensive transcriptional and translational reprogramming, leading to dynamic shifts in gene expression profiles. While numerous transcriptomic studies have characterized these responses across diverse plant species [4,5], proteomic level changes remain comparatively underexplored. Because host-pest interactions ultimately reshape the proteome through the expression of various defense-related genes, a proteomic perspective is crucial for identifying key proteins underlying herbivory induced responses and for deepening our understanding of plant-insect interactions.

Pigeon pea (*C. cajan*) is a major grain legume grown extensively in India. However, its production is severely constrained by the pod borer *H. armigera*, one of the most destructive pests globally. The pest’s diverse host range, capacity to inflict damage during sensitive crop stages such as flowering and pod formation, rapid life cycle, high dispersal ability, and propensity to evolve resistance against chemical insecticides make it exceptionally difficult to manage [6]. Although proteomics has been widely applied to investigate legume responses to abiotic stresses [7,8], pigeon pea defense responses to *H. armigera* have been investigated primarily at the genomic and transcriptomic levels our group [9,10] and other laboratories [11]. Similarly, our group has also studied changes in the pod wall proteome profile of chickpea due to *Helicoverapa armigera* attack [12]. Multi-omics studies in other crop systems have provided valuable insights into plant-insect interactions; for example, transcriptome-metabolome analyses have uncovered secondary metabolite reprogramming in cotton upon infestation by *Helicoverpa* and *Spodoptera* [13], while similar approaches in maize (*Spodoptera exigua*) and rice (*Cnaphalocrocis medinalis*) revealed key defense-responsive genes and pathways [14,15]. Extending such studies to pigeon pea at the proteome level would provide novel insights into herbivory-induced molecular responses, thereby advancing sustainable pest management strategies.

Mass spectrometry (MS)-based proteomics offers a robust platform for elucidating proteome alterations induced by biotic stresses. Our earlier work using 2D-PAGE coupled with MS identified DAPs in chickpea pod wall treated with *H. armigera* OS, detecting 35 and 20 DAPs in the commercial cultivar JG 11 and resistant cultivar ICC 506-EB, respectively, after 12 hours of simulated herbivory [12]. Similarly, Rathinam et al. [16] reported substantial proteomic reprogramming in pigeon pea (*C. cajan*) and its wild relative (*C. platycarpus*) at multiple time points post-infestation, although their experimental design involved direct larval feeding, which could introduce variability due to uncontrolled mechanical damage, larval movement, feeding heterogeneity, and frass deposition. A subsequent study employed Tandem Mass Tag (TMT)-based proteomics to compare proteome profiles of the wild-type resistant accession IBS 34719 and susceptible cultivar ICPL 87, revealing distinct proteomic signatures between the two genotypes [17]. However, this work did not include insect infestation or OS treatment, and the use of wild relatives with considerable genotypic and phenotypic divergence from cultivated germplasm complicates direct extrapolation of the results to pigeon pea breeding programs.

Earlier studies demonstrated that pigeon pea genotype ICPL 332 exhibits antixenosis-based resistance to *H. armigera*, being less preferred for oviposition compared to the susceptible genotype ICPL 87 [18]. ICPL 332 has been classified as moderately resistant, whereas ICPL 87 is highly susceptible to *H. armigera* infestation. To further elucidate the molecular basis of these contrasting phenotypes, we investigated the leaf proteome profiles of ICPL 332 and ICPL 87 following the application of *H. armigera* OS as a proxy for herbivory. This comparative proteomic approach aims to identify herbivory-responsive proteins and provide mechanistic insights into defense strategies deployed by pigeon pea against *H. armigera*.

## Materials and methods

### Plant material

Two pigeon pea (*C. cajan*) genotypes, ICPL 332 (moderately resistant) and ICPL 87 (susceptible), were procured from the International Crops Research Institute for Semi-Arid Tropics (ICRISAT), Patencharu, Hyderabad, India. The growth conditions of pigeon pea plants were maintained according to S. et al. [19]. For protein extraction, fresh leaves were collected from five-week-old plants, 12 h post application of *H. armigera* OS.

### Insect rearing, collection of OS, and plant treatment

The first-instar larvae of *H. armigera* were procured from ICRISAT-Hyderabad and reared at 25°C in insect-rearing trays on artificial diet composed of chickpea (37.5%), ascorbic acid (0.58%), sorbic acid (0.375%), aureomycin powder (1.43%), one multivitamin tablet, instant dry yeast (6%), and Difco agar (2.18%). After 2-3 weeks, pupae were transferred to plastic containers lined with absorbent cotton wool. Emerging adults were maintained in a 1:3 male-to-female ratio for mating, with container openings covered with cotton mesh for ventilation. Adults were fed with 25% sucrose solution adsorbed onto cotton wool. Eggs laid by females were allowed to hatch, and first-instar larvae were transferred to fresh rearing trays.

For collection of OS, mature larvae were gently inverted over microcentrifuge tubes, and the abdomen was stroked to release OS. The collected OS was stored at -80°C until further use. Simulated herbivory was performed by mechanically wounding leaves with a paper punch, followed by application of diluted OS (1:4; v/v) using a fine paint brush. Leaf tissues were collected 12 h post-treatment (hpt), snap-frozen in liquid nitrogen, and used for protein extraction.

### Detection of hydrogen peroxide

Hydrogen peroxide (H_2_O_2_) accumulation in response to simulated herbivory was quantified following the protocol of Labudda et al. [20] with minor modifications. Approximately 100 mg of leaf tissue was homogenized with 1 mL of an extraction mixture containing 10 mM sodium phosphate buffer (pH 5.8), 1 M potassium iodide, and 0.1% trichloroacetic acid (1:2:1; v/v/v). The homogenates were centrifuged at 16,000×g for 15 min at 4°C, and the resulting supernatants were incubated in the dark for 20 min at room temperature. Samples were again centrifuged at 16,000×g for 10 min, and the absorbance was recorded at 350 nm using a Basic BioSpectrometer (Eppendorf). H_2_O_2_ concentrations were determined from a standard curve and expressed as mM g^-1^ fresh weight.

In addition to quantitative estimation, qualitative detection of H_2_O_2_ was performed using the 3,3′-diaminobenzidine (DAB) staining method described by Liu and Tim [21]. A 1 mg mL^-1^ DAB solution (pH 3.6) was prepared by dissolving the reagent in distilled water for overnight at 37°C with continuous shaking at 200 rpm. Leaf samples, harvested 12 h post-treatment, were immersed in DAB solution and incubated in the dark at 37°C with shaking (130 rpm) for 16 h. Following incubation, the solution was replaced with ethanol:acetic acid (3:1; v/v) and samples were boiled in a water bath at 100°C for 1-2 h to remove chlorophyll pigments. Finally, tissues were cleared by boiling in lacto-glycerol solution (lactic acid:glycerol:H_2_O; 1:1:1; v/v/v) for 1 h at 100°C. The stained leaves were imaged and used for subsequent qualitative analysis.

#### Antioxidant enzymes assays

To evaluate the activity of key antioxidant enzymes, including peroxidase (POD), superoxide dismutase (SOD), and catalase (CAT), in response to simulated herbivory, biochemical assays were performed. Approximately 100 mg leaf tissue was homogenized in liquid nitrogen and resuspended in 3 mL of extraction buffer containing 0.1 M phosphate buffer (pH 7.5) and 0.5 mM EDTA. The homogenate was filtered and centrifuged at 15,000 × g for 20 min, and the resulting supernatant was used as the enzyme extract. Enzymatic activity of POD, SOD, and CAT were determined following protocols of Castillo et al. [22], Dhindsa et al. [23], and Aebi [24], respectively, with minor modifications.

#### Total protein extraction for proteomic analysis

Total protein was extracted from 1 g leaf tissue collected from untreated (control) and treated plants following the protocol of S. et al. [19]. Leaf samples were collected in triplicate from both groups to ensure reproducibility. The isolated protein pellet was resuspended in 200 µl of the ReadyPrep™ 2-D starter kit Rehydration/sample buffer (Bio-Rad, USA). Protein concentration was determined using Bradford assay, and the samples were subsequently used for two-dimensional SDS-PAGE analysis.

#### Two-dimensional SDS-PAGE (2D-PAGE) analysis and MS analysis

Proteins were separated using two-dimensional SDS-PAGE (2D-PAGE). Immobilized pH gradient (IPG) strips (pH 4–7, 7 cm; ReadyStrip™ IPG Strips, Bio-Rad, USA) were passively rehydrated and loaded with approximately 50 µg proteins. Isoelectric focusing (IEF) was performed using the PROTEAN®i12™ IEF system (Bio-Rad, USA) using the manufacturer’s protocol (Table 1). The IEF, equilibration, SDS-PAGE, silver staining and MS conditions were followed according to S. et al. [19].

**Table 1.** PROTEAN®i12™ IEF system condition for IEF.

| Step | Voltage | Gradient | Current | Value | Units |
| --- | --- | --- | --- | --- | --- |
| 1 | 250 | Linear | 50 | 0:20 | HH:MM |
| 2 | 4000 | Linear | 50 | 2:00 | HH:MM |
| 3 | 4000 | Rapid | 50 | 10,000 | Volt Hr |

For MS analysis, 40 DAPs reproducibly detected across three biological replicates from both cultivated pigeon pea genotypes were excised and subjected to proteomic profiling. Peptides were analyzed using an Easy-nLC 1000 system coupled to an Orbitrap Exploris mass spectrometer. Raw spectra data were processed using Proteome Discoverer (v2.5) software against the UniProt *Cajanus cajan* protein database [25]. Candidate proteins were selected based on peptide-spectrum match (PSM) scores. Protein identification was performed using Sequest and Amanda search algorithms, with precursor and fragment mass tolerances of 10 ppm and 0.02 Da, respectively. Protein identifications were filtered at a stringent false discovery rate (FDR) of 1% (q ≤ 0.01). Relative fold changes (FCs) between the two genotypes were quantified in triplicate using ImageJ. In addition, a heatmap was generated for the 10 common DAPs identified in both genotypes using the R statistical software (R Core Team, Austria).

#### RNA extraction, cDNA synthesis and quantitative PCR analysis

Total RNA extracted from leaves of five-week-old pigeon pea plants at 12 h after post-treatment using the RNeasy plant mini kit (Qiagen, Germany), following the protocol described by Meshram et al. [10]. Control and treated samples were collected in biological triplicates. For quantitative PCR (qPCR), 30 ng of cDNA was used as a template in reactions prepared with PowerUp™ SYBR® Green Master Mix (Applied Biosystems, USA) and performed on the QuantStudio™ 5 Real time PCR system (Applied Biosystems, USA), according to the manufacturer’s protocol. Gene specific primers were designed using the OligoPerfect™ Designer tool (https://www.thermofisher.com/apps/oligoperfect/design) (Table S1). Expression levels were normalized using *Tubulin* as an endogenous reference gene.

#### Statistical analysis

The experimental data from biochemical analysis and qPCR analysis were statistically analyzed using a two-tailed Student’s t-test with two sample groups assuming equal variances. The corrected p-value < 0.05 was considered significant. The graphical representations of the data were done with GraphPad Prism software (https://www.graphpad.com).

## Results and Discussion

### Application of OS induces reactive oxygen species (ROS) generation in pigeon pea

Under stress conditions, plants generate ROS, which act as key indicators of host-pathogen interactions and the activation of downstream signaling pathways. Kuźniak and Kopczewski [26] reported that elevated ROS production within chloroplasts is associated with a decline in photosynthetic rate. To assess ROS accumulation in cultivated pigeon pea genotypes, we performed an H_2_O_2_ assay at 12-, 24- and 48-h post simulated herbivory (Fig. 1a). A significantly higher level of H₂O₂ was detected in treated leaf tissues of both genotypes compared to control plants at 12 h and 24 h. This regulated ROS generation plays a pivotal role in activating defense mechanisms against pathogen attack and contributes to the modulation of the overall stress response in plants. The maximum accumulation of H₂O₂ was observed at 12 h post-treatment in both genotypes, followed by a gradual decline, with the lowest levels recorded at 48 h irrespective of genotype. Based on this accumulation trend, the 12 h time point was selected for subsequent proteomic analysis in response to simulated herbivory.

**Fig.1.**
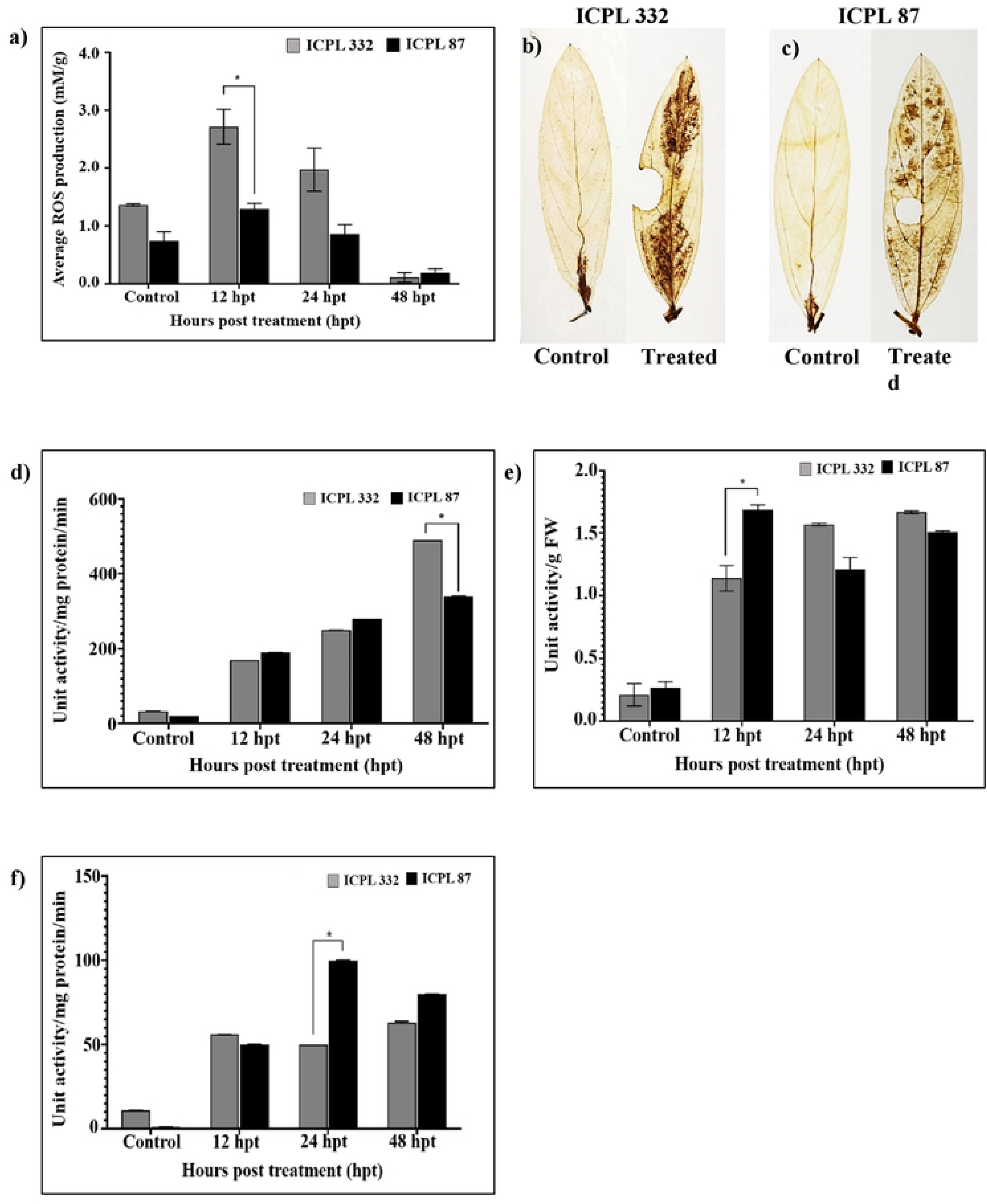

Furthermore, a DAB assay was performed to qualitatively visualize ROS accumulation at 12 h post-simulated herbivory (Fig. 1b, 1c). In agreement with the quantitative assay, significantly elevated H₂O₂ levels were observed in treated leaf samples of both genotypes compared to controls. Notably, H₂O₂ accumulation was higher in the moderately resistant cultivar ICPL 332 than in the susceptible ICPL 87 at 12 h post-treatment. This finding is consistent with earlier reports of ROS accumulation in ICPL 332 following *Helicoverpa* infestation [9,10], indicating that simulated herbivory induces ROS production, which in turn activates defense signaling pathways and contributes to both localized and systemic responses against potential herbivore challenges.

### Enhanced ROS levels trigger antioxidant defense mechanisms in pigeon pea

Stressful environmental conditions disturbs cellular homeostasis and ion balance in plants, leading to osmotic stress that stimulates the overproduction of ROS. Excessive accumulation of ROS results in damage to cellular organelles and membranes via lipid peroxidation, degradation of vital biomolecules, and in severe cases, cell death. To counteract this, plants rely on antioxidant enzymes, which constitute the first line of defense against ROS-induced oxidative injury and are indispensable for maintaining redox equilibrium [27]. Evidence from earlier studies demonstrates that induction of enzymatic and non-enzymatic antioxidants is a critical adaptive response, enabling cells to overcome oxidative damage imposed by environmental stresses [28].

In the present study, we evaluated the activity of key antioxidant enzymes - POD (Fig. 1d), SOD (Fig. 1e) and CAT (Fig. 1f) - in cultivated pigeon pea genotypes subjected to simulated herbivory, at 12-, 24-, and 48-h post treatment. Across all time intervals, both genotypes exhibited higher antioxidant activity relative to untreated controls. Notably, ICPL 87 showed markedly elevated antioxidant enzyme activity at 12 h compared to ICPL 332. Although ICPL 332 displayed relatively lower antioxidant activity at 12 h, it showed a progressive increase at 48 h, a pattern not evident in ICPL 87. The reduced antioxidant enzyme activity in ICPL 87 may be resulted due to the downregulation of the P-type Cu^+^ transporter (COPT; A0A151TJQ8; Spot No. S17) and glutaredoxin-dependent peroxiredoxin (GPx-Q; A0A151TVB0; Spot No. S18), as identified in our proteomic dataset. COPT has been established as a central regulator of copper homeostasis, directly influencing the enzymatic activity of Cu/Zn superoxide dismutase (Cu/ZnSOD), a pivotal antioxidant enzyme [29]. Likewise, GPx-Q plays an essential role in peroxide detoxification under oxidative stress [30]. Our results are in consistent with prior reports showing that enhanced antioxidant activity contributes to the activation of plant defense systems and facilitates tolerance to diverse abiotic stresses. Furthermore, we observed that elevated ROS levels at the 12 h coincided with suppressed antioxidant enzyme activity, whereas later time points revealed an inverse trend, where enhanced antioxidant activity corresponded to reduced ROS accumulation.

### Identification of DAPs in pigeon pea in response to simulated herbivory

To investigate the signaling mechanisms operating during the interaction between pigeon pea (*C. cajan*) and *H. armigera*, we examined the leaf proteome of two cultivated pigeon pea genotypes 12 h after simulated herbivory. Total proteins were extracted from control and treated samples in triplicate and subjected to 2D-PAGE analysis. Approximately 220 protein spots were detected in ICPL 332, whereas about 200 spots were observed in ICPL 87 (Fig. 2). Image analysis revealed ∼40 DAPs in the susceptible cultivar ICPL 87 and ∼35 DAPs in the moderately tolerant cultivar ICPL 332. From the 75 DAPs across both genotypes, 40 reproducible spots (21 from ICPL 87 and 19 from ICPL 332) were selected for MS analysis, based on consistent detection in SDS-PAGE profiles obtained from triplicates samples to reduce error and misinterpretation (Fig. S1, S2). Among these 40 DAPs, 10 (hereafter referred to as common proteins) were present in both genotypes and further used for comparative statistical analysis (Fig. 3). Fold change (log2 fold) of the DAPs were quantified using the ImageJ software to assess the differential proteomic responses under simulated herbivory by OS of *H. armigera*.

**Fig.2.**
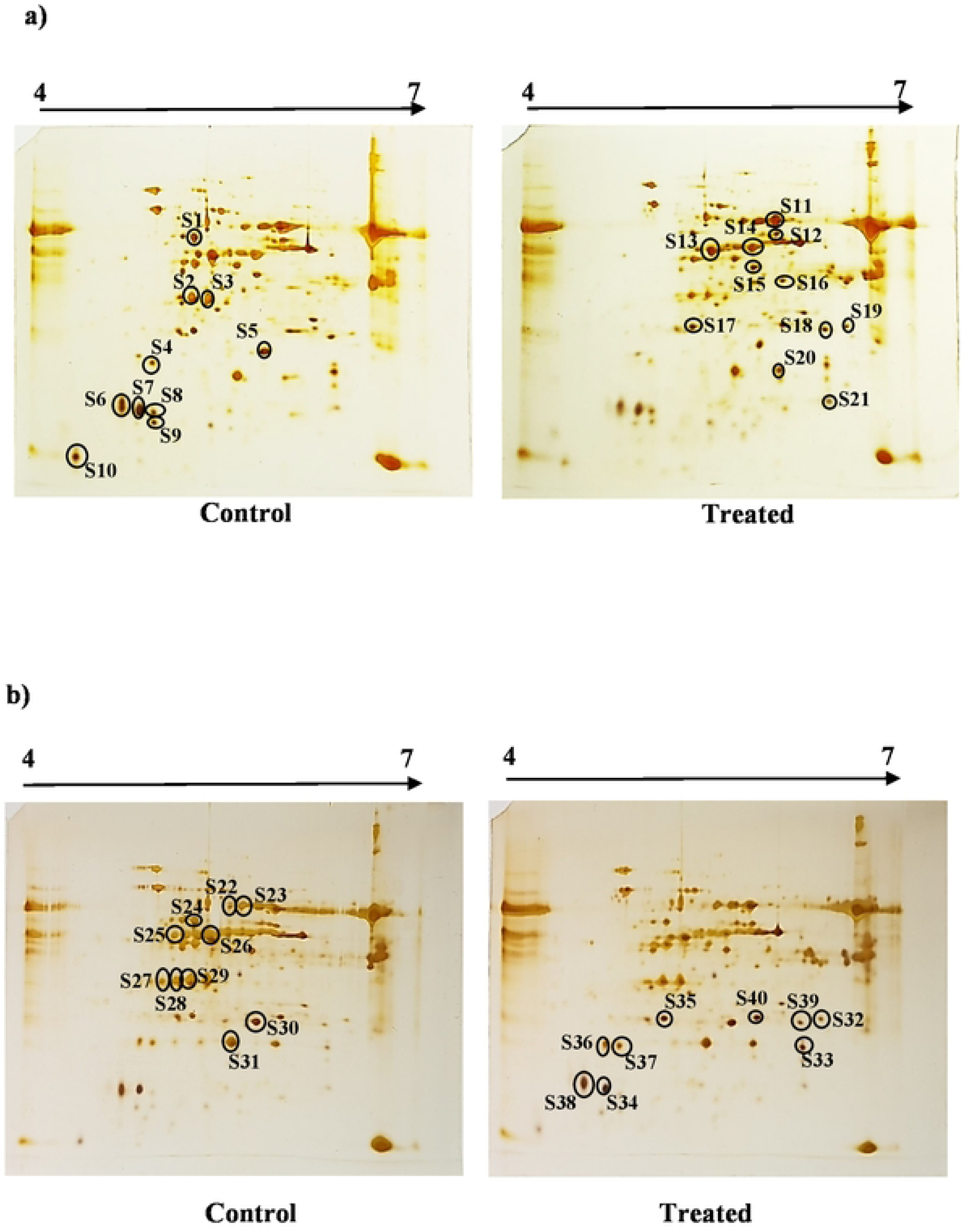

**Fig.3.**
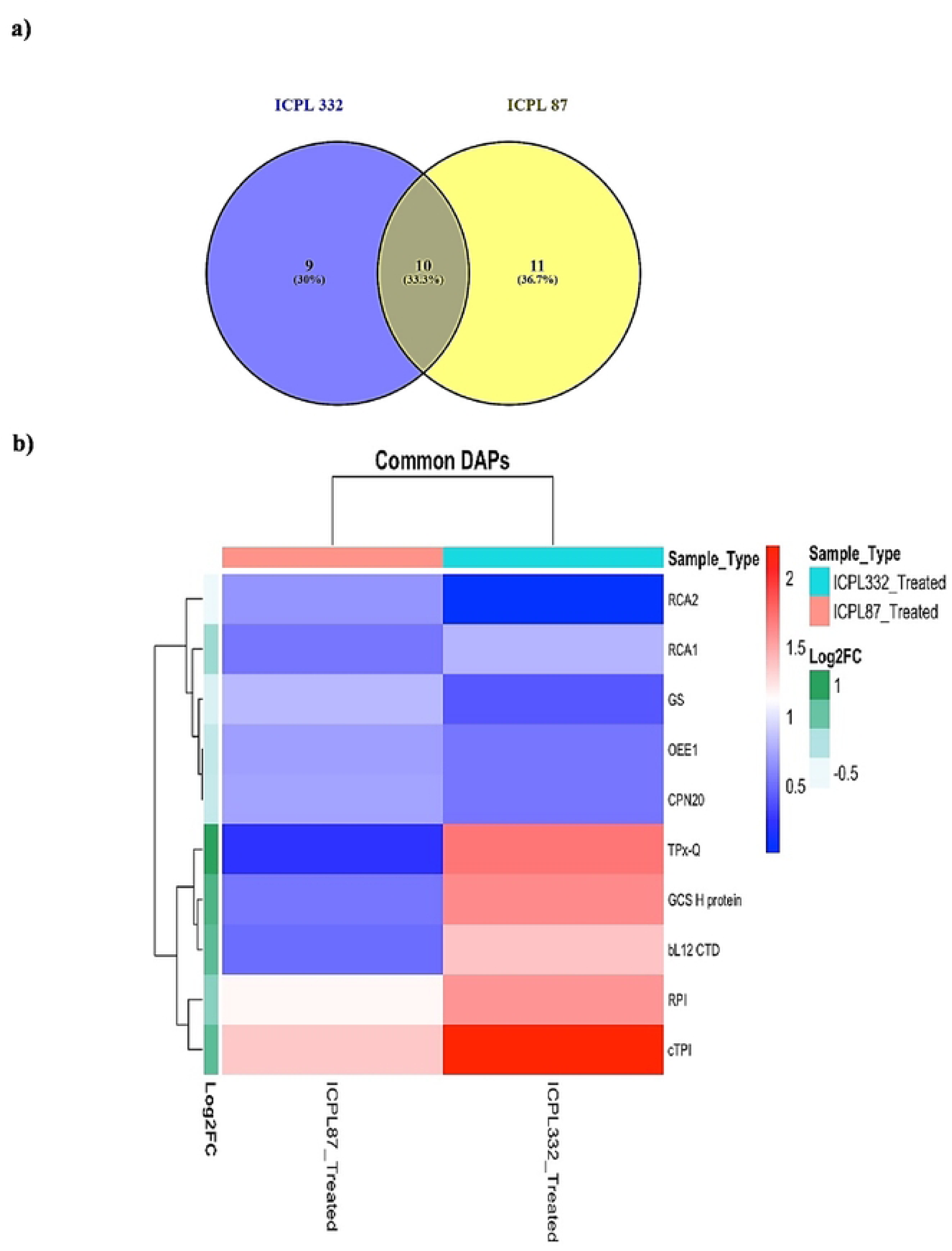

The identified DAPs were classified into three functional categories: (a) primary metabolism, (b) stress response, and (c) growth and development. Of the 21 DAPs identified in ICPL 87, 10 were common proteins (six related to primary metabolism, three to stress responses and one to growth and development; Fig. 4a, 4b; Table 2), whereas the remaining 11 were unique (six associated with primary metabolism, two with stress response, one with growth and development and two with an unknown function; Fig 5a, 5b; Table 3). Similarly, among the 19 DAPs identified in ICPL 332, nine were unique (five involved in primary metabolism, three in stress response and one of unknown function; Fig 5c, 5d; Table 4), while 10 were common proteins shared with ICPL 87 (six involved in primary metabolism, three in stress responses and one in growth and development; Table 2).

**Fig.4.**
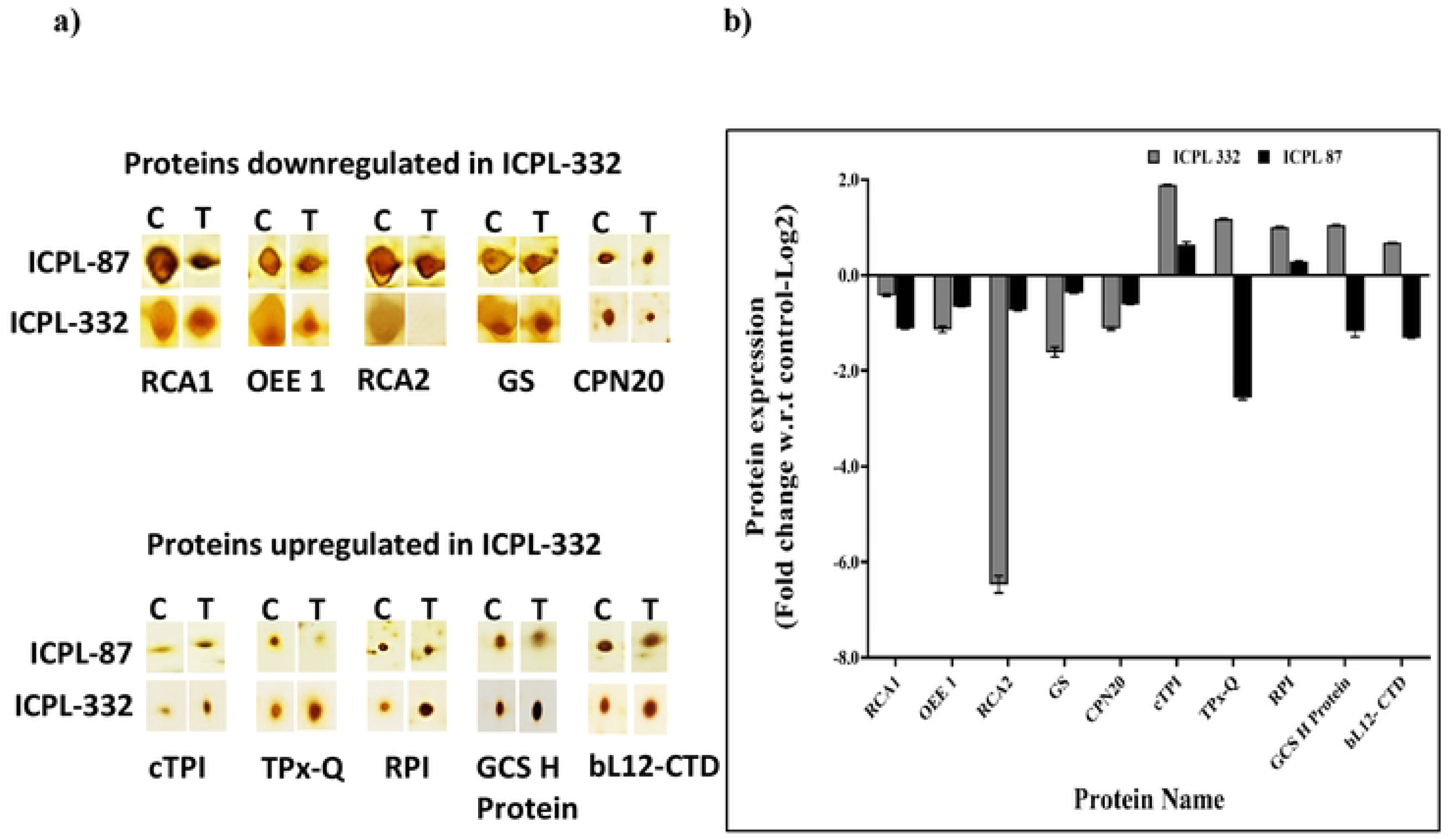

**Fig.5.**
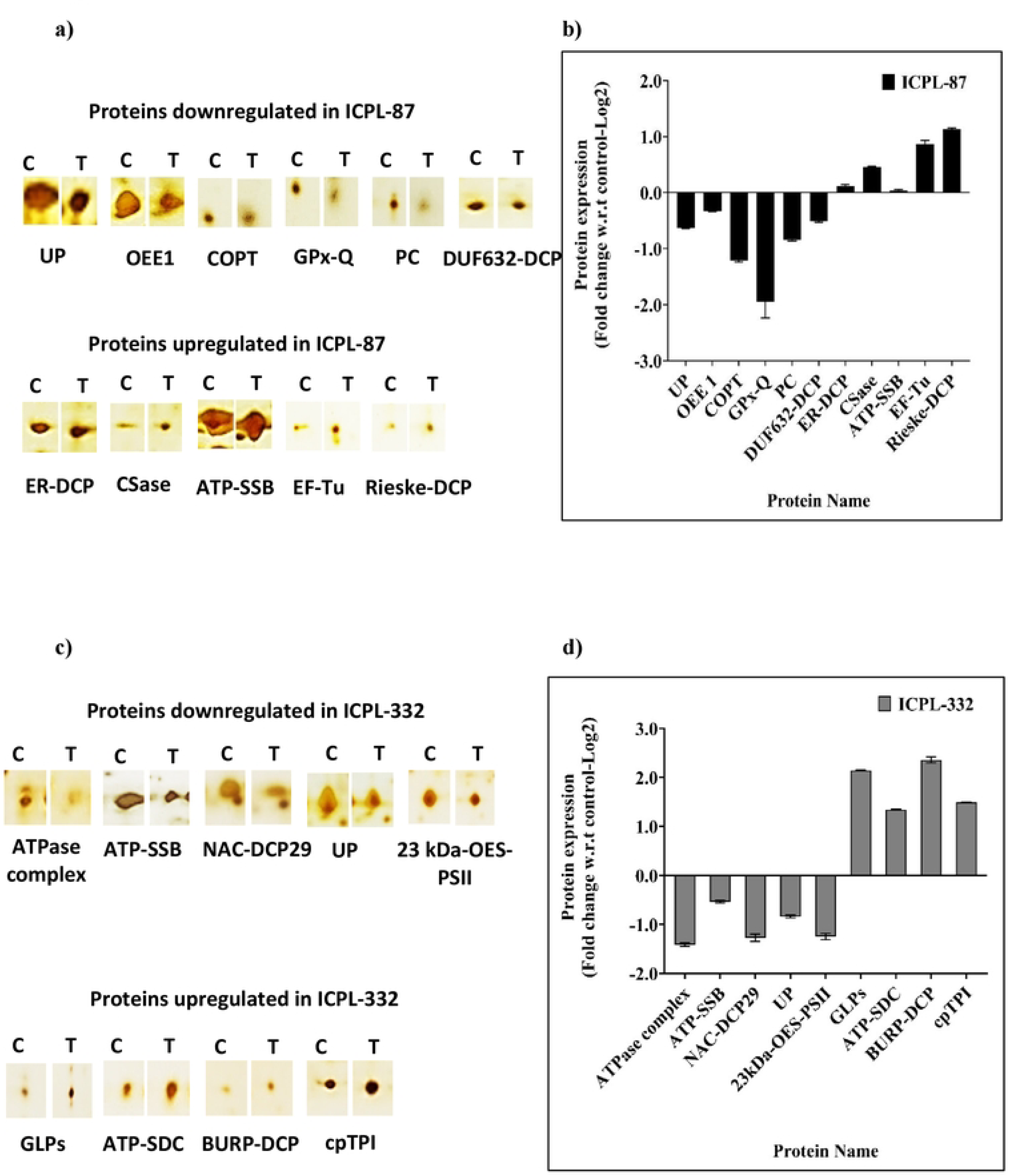

**Table 2:**
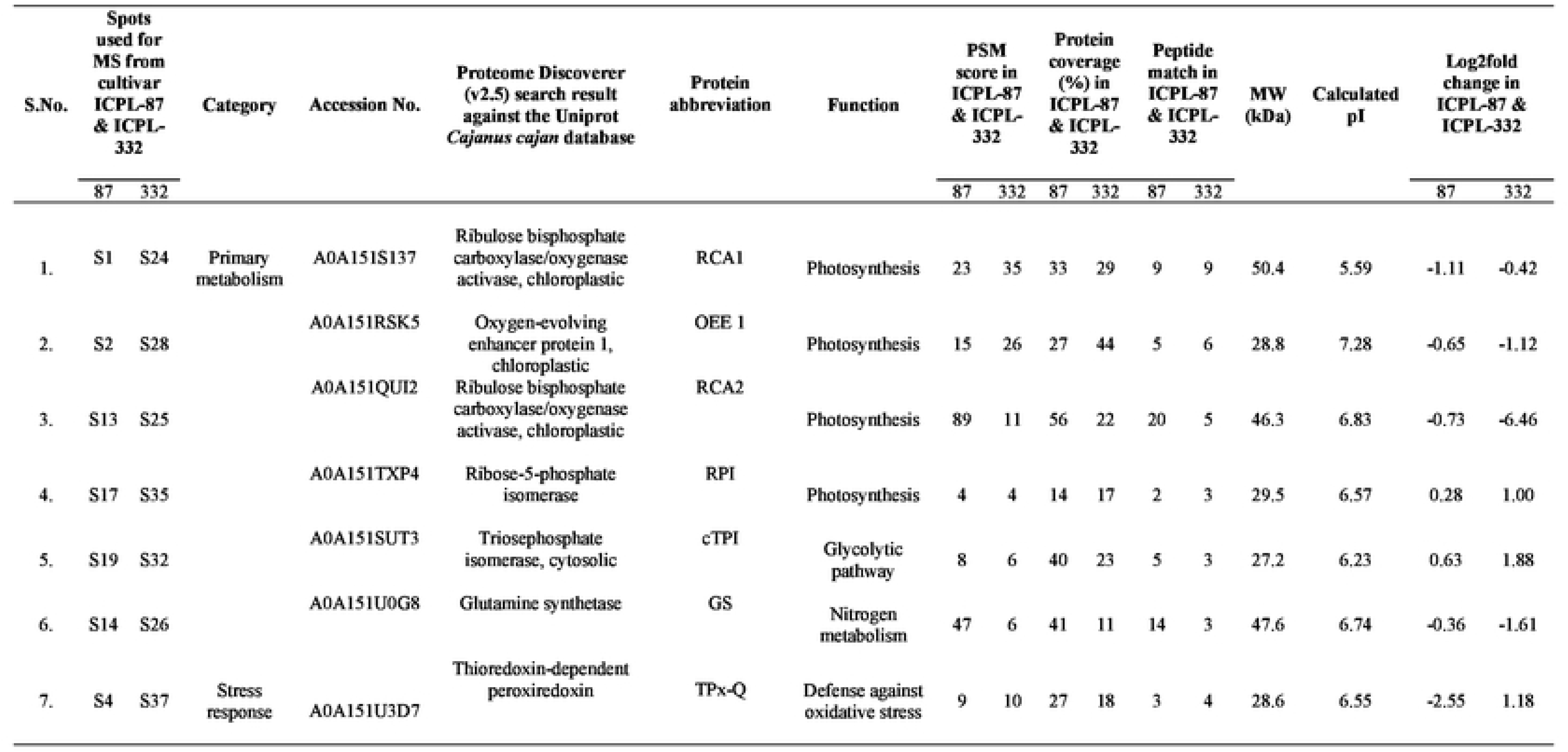
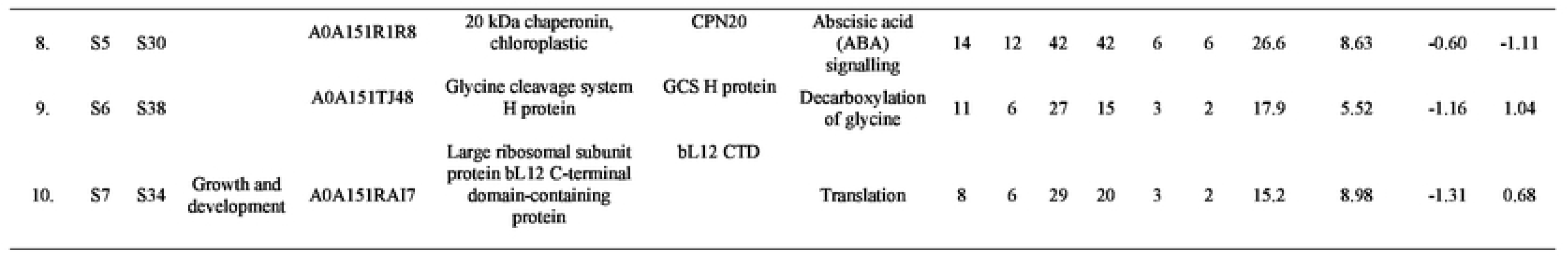
List of common proteins identified in Pigeon pea (ICPL-87 and ICPL-332) 12 h post treatment.

**Table 3:**
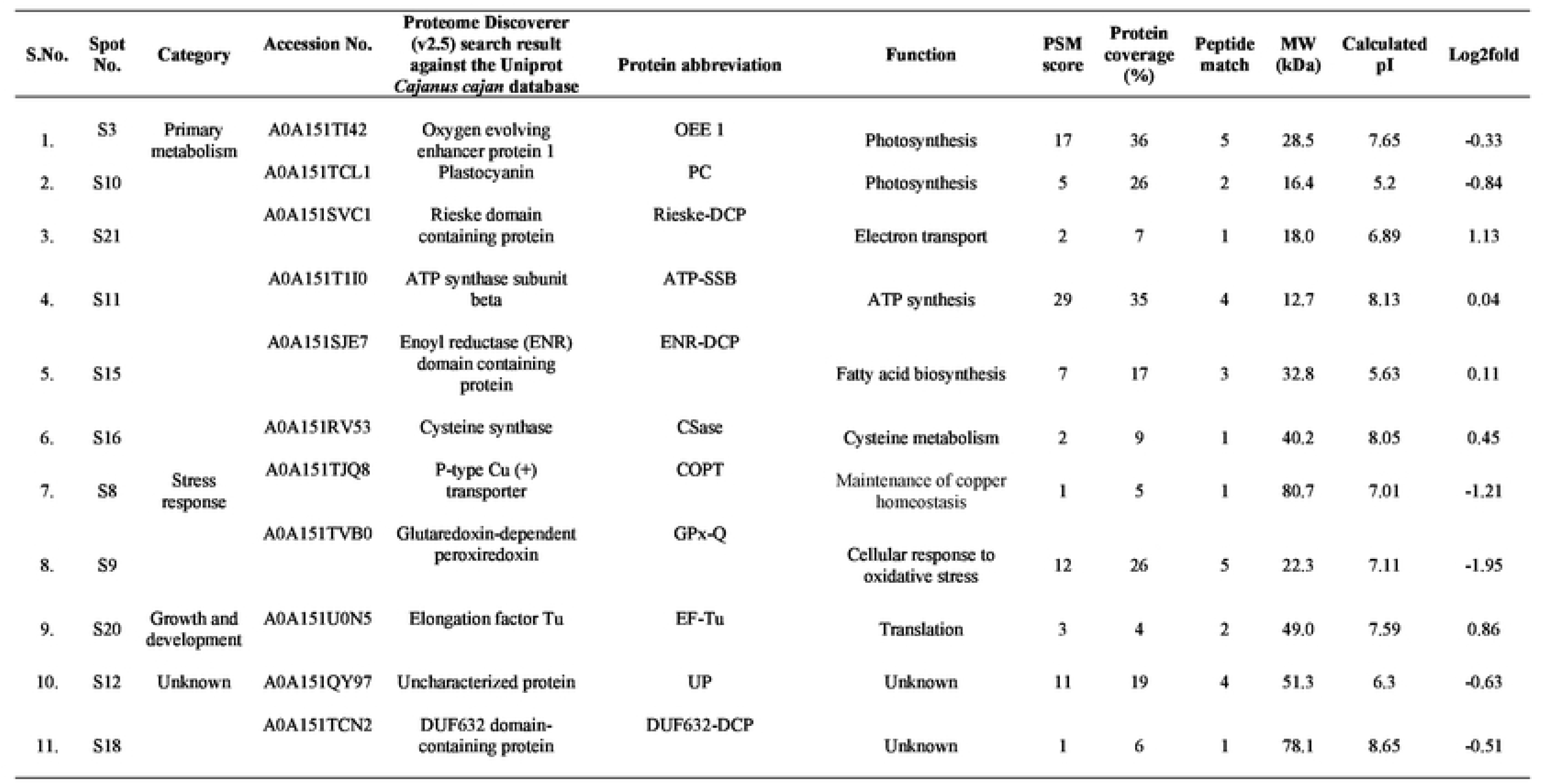
List of identified proteins in Pigeon pea (JCPL-87) 12 h post treatment.

**Table 4:** List of identified proteins in Pigeon pea (JCPL-332) 12 h post treatment.

| S.No. | Spot No. | Category | Accession No. | Proteome Discoverer (v2.5) search result against the Uniprot *Cajanus cajan* database | Protein abbreviation | Function | PSM score | Protein coverage (%) | Peptide match | MW (kDa) | Calculated pI | Log2fold |
| --- | --- | --- | --- | --- | --- | --- | --- | --- | --- | --- | --- | --- |
| 1. | S31 | Primary metabolism | A0A151RKE1 | 23 kDa subunit of oxygen evolving system of photosystem II | 23kDa-OES-PSII | Photosynthesis | 24 | 44 | 7 | 24.4 | 6.93 | -1.24 |
| 2. | S22 |  | A0A151TCV6 | ATPase F1/V1/A1 complex alpha/beta subunit N terminal domain containing protein | ATPase complex | ATP metabolic process | 4 | 24 | 1 | 6.9 | 9.99 | -1.41 |
| 3. | S23 |  | A0A151TII0 | ATP Synthase subunit beta | ATP-SSB | ATP synthesis | 14 | 28 | 2 | 12.7 | 8.13 | -0.53 |
| 4. | S36 |  | A0A151U827 | ATP synthase delta chain, chloroplastic | ATP-SDC | ATP synthesis | 11 | 20 | 4 | 27.1 | 6.96 | 1.34 |
| 5. | S40 |  | A0A151TVE0 | Triose phosphate isomerase, chloroplastic | CpTPI | Calvin-Benson cycle | 9 | 19 | 4 | 32.9 | 7.30 | 1.49 |
| 6. | S33 | Stress response | A0A151RWD4 | Germin-like protein | GLPs | Biotic/abiotic stress responsive protein | 3 | 19 | 2 | 23.2 | 6.76 | 2.14 |
| 7. | S27 |  | A0A151T9N7 | NAC domain containing protein 29 | NAC-DCP29 | Transcription regulation | 1 | 11 | 1 | 38.2 | 9.00 | -1.27 |
| 8. | S39 |  | A0A151U7C1 | BURP domain containing protein 3 | BURP-DCP | Stress responsive | 1 | 16 | 1 | 26.6 | 7.12 | 2.35 |
| 9. | S29 | Unknown | A0A151QY97 | Uncharacterized protein | UP | Unknown | 16 | 25 | 8 | 51.3 | 6.30 | -0.83 |

### Altered photosynthetic rate in pigeon pea cultivated genotypes post simulated herbivory

Photosynthesis in plants is markedly influenced under biotic stress, such as pathogen attack or insect herbivory. During such conditions, plants frequently reallocate energy and metabolic resources from photosynthesis and growth toward defense responses, including the production of ROS and defense-related proteins. These metabolic adjustments collectively suppress photosynthetic efficiency, limit biomass accumulation, and, under sustained or severe stress, compromise plant productivity and yield.

To assess the influence of simulated herbivory-induced stress response on pigeon pea, we first examined the differential regulation of photosynthesis- and stress-related proteins. MS data revealed that, among the commonly expressed proteins, several key photosynthesis-related proteins, including oxygen-evolving enhancer protein 1 (OEE1; A0A151RSK5; Spot No. S2; S23), ribose-5-phosphate isomerase (RPI; A0A151TXP4; Spot No. S4; S25), and two isoforms of ribulose bisphosphate carboxylase/oxygenase (RuBisCO) activases (RCA1; A0A151S137; Spot No. S1; S22 and RCA2; A0A151QUI2; Spot No. S3; S24), were differentially regulated in both cultivated genotypes following simulated herbivory. In addition, two unique photosynthesis-related DAPs were detected in ICPL 87, namely plastocyanin (PC; A0A151TCL1; Spot No. S12) and an isoform of OEE1 (A0A151TI42; Spot No. S11), whereas one unique protein, the 23 kDa subunit of the oxygen evolving system of photosystem II (23kDa-OES-PSII; A0A151RKE1; Spot No. S32), was observed in ICPL 332.

Among these proteins, RuBisCO activase (RCA) plays a critical role in RuBisCO activation but is known to undergo downregulation under stress conditions. In *Arabidopsis*, elevated jasmonic acid (JA) levels under stress has been shown to repress RCA through F-box protein coronatine-insensitive 1 (COI1)-mediated degradation [31]. Consistent with these findings, RCA was significantly downregulated in both pigeon pea genotypes under simulated herbivory.

Similarly, OEE1, an essential component of photosystem II, was also downregulated in both genotypes. Although its role under abiotic stress, including drought, salinity, and oxidative stress, has been associated with chloroplast degradation through interactions with PsbO1 [32], its function under biotic stress remains less clearly defined. Previous studies suggest that in addition to regulating light-driven water splitting and maintaining photosynthetic electron transport (PET), OEE1 possesses H_2_O_2_ scavenging activity [33]. Therefore, downregulation of OEE1 impairs PET, lowers photosynthetic efficiency and promotes ROS accumulation. In line with this, our study demonstrated significant OEE1 downregulation in both genotypes, with a significant effect in ICPL 332.

Furthermore, PC, an electron carrier in photosynthesis, was found to be downregulated in ICPL 87. Given its central role in PET, suppression of PC disrupts electron transport and enhances ROS accumulation [34]. Likewise, in ICPL 332, the 23 kDa-OES-PSII, a protein essential for photosynthesis and oxygen evolution [35], was significantly downregulated. Earlier reports indicate that loss of this protein reduces O2-evolving activity by lowering the affinity of the oxygen-evolving complex (OEC) for chloride ions, thereby leading to H_2_O_2_ accumulation [36]. Taken together, our results clearly demonstrate altered photosynthetic performance in pigeon pea under simulated herbivory. This disruption of photosynthetic processes perturbs electron transport and energy flow within chloroplasts, triggering ROS production [26], which was further supported by our biochemical assays.

#### ROS-Mediated signaling influences stress related proteins under simulated herbivory

ROS function as key signaling molecules during environmental stress by modulating downstream transcriptional regulation through the activation of various transcription factors [37]. In the present study, we observed that simulated herbivory triggered differential regulation of several stress-associated proteins and components of the translational machinery in pigeon pea cultivated genotypes. Among the identified proteins, chloroplast-localized chaperonin 20 kDa (CPN20; A0A151R1R8; Spot No. S8; S29), thioredoxin-dependent peroxiredoxin (TPx-Q; A0A151U3D7; Spot No. S7; S28), and glycine cleavage system H protein (GCS H; A0A151TJ48; Spot No. S9; S30) exhibited genotype-specific modulation. TPx-Q, which plays a crucial role in protecting the photosynthetic machinery under oxidative stress [38], was significantly upregulated in ICPL 332 but downregulated in ICPL 87, suggesting a stronger oxidative stress response in ICPL 332. Similarly, GCS H, involved in amino acid metabolism and photorespiratory carbon flux [39], showed upregulation in ICPL 332 and downregulation in ICPL 87. Consistent with previous findings in tobacco, where GCS H overexpression enhanced stress adaptation, biomass production, and photosynthetic efficiency [40], our results indicate that ICPL 332 employs a more effective adaptive strategy. In contrast, CPN20, implicated in protein folding and abscisic acid (ABA) signaling [41], was downregulated in both genotypes, indicating possible suppression of its role under herbivory-induced stress.

In addition, ICPL 332 displayed increased accumulation of several stress-responsive proteins, including germin-like proteins (GLPs; A0A151RWD4; Spot No. S37), BURP domain-containing protein 3 (BURP DCP 3; A0A151U7C1; Spot No. 39), and NAC domain-containing protein 29 (NAC DCP 29; A0A151T9N7; Spot No. 38). GLPs and BURP DCP 3 are known to contribute stress tolerance [42], while BURP family genes in soybean have been reported to respond to stress owing to stress-related cis-elements present in their promoter regions [43]. Interestingly, despite their importance, NAC DCP 29, a well-characterized transcription factor involved in stress regulation [44], was downregulated, pointing toward a complex transcriptional control in ICPL 332 under herbivory stress.

In the case of translational machinery, the large ribosomal subunit protein bL12 C-terminal domain-containing protein (bL12 CTD; A0A151RAI7; Spot No. S10; S31) showed genotype-specific expression pattern. This chloroplast-associated 50S ribosomal protein [25] contains a C-terminal domain that facilitates interaction with translation factors. Although its precise stress-related role remains unclear, bL12 transcripts have been reported to increase in response to heat [45] and high-intensity light stress [46], likely as a compensatory mechanism to maintain photosynthetic efficiency. Under simulated herbivory, however, bL12 was significantly downregulated in ICPL 87 but strongly upregulated in ICPL 332, reinforcing the latter’s superior stress tolerance. Furthermore, elongation factor Tu (EF-Tu; A0A151U0N5; Spot No. S19), which mediates GTP-dependent aminoacyl-tRNA binding during translation, was slightly upregulated in ICPL 87. While EF-Tu has been implicated in preventing protein aggregation under stress [47], its limited accumulation in ICPL 87 suggests comparatively weaker stress adaptation than in ICPL 332.

Taken together, these proteomic insights highlight that ICPL 332 possesses a more effective defense response to simulated herbivory than ICPL 87, as reflected by the upregulation of critical stress-related proteins and components of the translational machinery. The genotype-specific regulation of stress-associated proteins and ribosomal machinery underscores distinct adaptation strategies, with ICPL 332 demonstrating enhanced resilience to herbivory-induced oxidative stress. Furthermore, the differential expression of numerous primary metabolism-related proteins across both genotypes suggests a coordinated response, where herbivory stress not only activates defense mechanisms but also reprograms core metabolic pathways to balance energy demands and sustain cellular functions during stress adaptation.

#### Quantitative PCR (qPCR) analysis of selected genes

To further corroborate the proteomics findings, qPCR was employed to assess the transcript abundance of six key common proteins that displayed differential expression pattern, thereby determining whether the observed variations at the protein level were also mirrored at the mRNA level following 12 h post-simulated herbivory (Fig. 6). In the cultivated genotype ICPL 332, genes associated with photosynthetic processes, specifically RCA1 (A0A151S137) and OEE1 (A0A151RSK5), demonstrated a clear downregulation at the transcript level, suggesting a suppression of photosynthetic activity under herbivore-induced stress. In contrast, the stress-responsive gene TPx-Q (A0A151U3D7) showed a slight but notable upregulation, which was in agreement with the corresponding protein-level data, thereby reinforcing its potential role in stress adaptation. On the other hand, in the cultivated genotype ICPL 87, a distinct expression profile was observed wherein all the selected genes, including TPx-Q, were consistently downregulated, with the only exception being a minor upregulation recorded in the RCA1 gene. This genotype-specific divergence in transcriptional regulation highlights the differential stress-response strategies employed by the two pigeon pea genotypes when exposed to herbivory.

**Fig.6.**
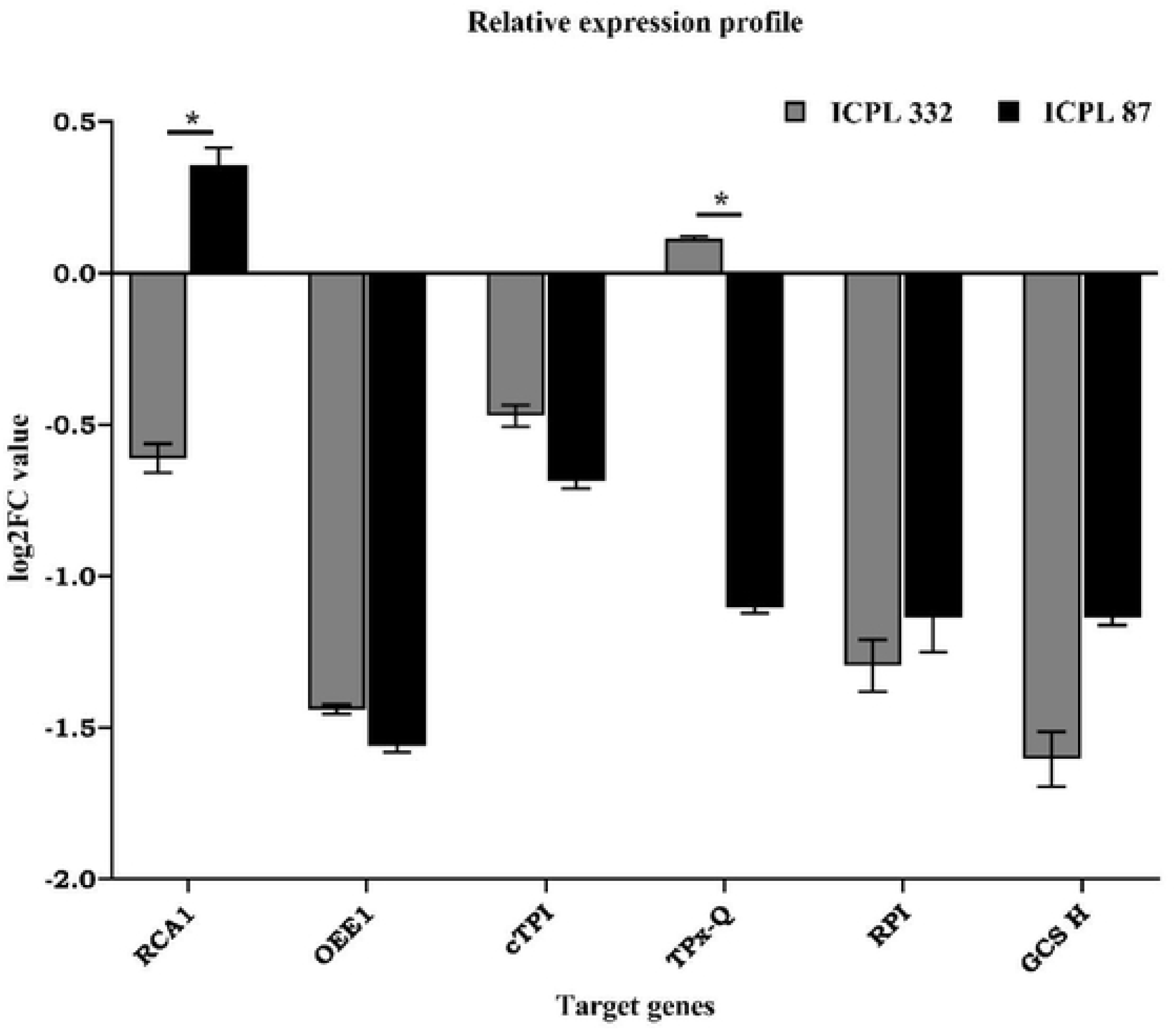

## Conclusion

This study provides valuable insights into the proteomic alterations occurring in pigeon pea leaves following simulated herbivory. The generation of ROS under simulated herbivory conditions triggered the activation of antioxidant enzymes and modulated stress-responsive proteins at the 12 h time point. Differential regulation of proteins associated with primary metabolism, stress adaptation, and growth-related processes underscores the complexity of plant defense strategies. Notably, the stronger downregulation of stress-responsive proteins in the susceptible cultivar (ICPL 87) relative to the moderately resistant cultivar (ICPL 332) indicates a genotype-dependent variation in defense responses. Collectively, these findings enhance our understanding of the molecular mechanisms underlying herbivory-induced stress adaptation in pigeon pea and hold promise for the development of pest-resistant cultivars through selective breeding and biotechnological interventions. Nevertheless, further investigations are needed to clarify the functional significance of the unidentified proteins and their potential contributions to stress tolerance.

## Author Contributions

S.A.: supervision, conceptualization, data curation, writing – review and editing, project related administration. B.K.S. Administration, A.S.: methodology, data analysis, writing of original draft. P.J.K.: data analysis, writing of original draft. S.K.M: methodology. J.J.: maintained the insect larvae for extraction of OS. A.D.: extracted the OS. R.I.P.: methodology. D.D.: writing – review and editing, S.D: methodology.

## Acknowledgment

AS acknowledges the financial support received through the Department of Science and Technology-Innovation in Science Pursuit for Inspired Research (DST-INSPIRE) Fellowship (Application No. DST/INSPIRE/03/2022/001942, Reg No. IF210021) from the Department of Science and Technology, Government of India, for the pursuit of her doctoral research.

## Ethics declarations

### Ethics approval and consent to participate

Not applicable

### Consent for publication

Not applicable.

### Approval of the research protocol by an institutional reviewer board

Not applicable.

### Informed consent

Not applicable.

### Registry and the registration no. of the study/trial

Not applicable.

### Animal studies

Not applicable.

### Clinical trial number

Not applicable.

